# Functional dynamics of dopamine synthesis during monetary reward and punishment processing

**DOI:** 10.1101/2019.12.23.886812

**Authors:** Andreas Hahn, Murray B. Reed, Verena Pichler, Paul Michenthaler, Lucas Rischka, Godber M. Godbersen, Wolfgang Wadsak, Marcus Hacker, Rupert Lanzenberger

## Abstract

**Purpose:** In the human brain endogenous dopamine release is commonly assessed by the PET competition model. Although thoroughly validated, cognitive processing yields low signal changes and the assessment of several task conditions requires repeated scanning. Using the framework of functional PET imaging we introduce a novel approach which leverages the incorporation of the radioligand 6-[^18^F]FDOPA into the dynamic fast-acting regulation of the corresponding enzyme activities by neuronal firing and neurotransmitter release. We demonstrate the feasibility of the approach by the assessment of widely described sex differences in dopamine neurotransmission.

**Methods:** Reward and punishment processing was behaviorally investigated in 36 healthy participants, where 16 underwent fPET and fMRI while performing the monetary incentive delay task. 6-[^18^F]FDOPA was applied as bolus+infusion during a single 50 min PET acquisition. Task-specific changes in dopamine synthesis were identified with the general linear model and quantified with the Gjedde-Patlak plot.

**Results:** Monetary gain induced 78% increase in nucleus accumbens dopamine synthesis vs. 49% for loss in men. Interestingly, the opposite was discovered in women (gain: 51%, loss: 78%). Behavioral modeling revealed direct associations of task-specific dopamine synthesis with reward sensitivity in men (rho = −0.7) and with punishment sensitivity in women (rho = 0.89). As expected, fMRI showed robust task-specific neuronal activation but no sex difference.

**Conclusions:** Our findings provide a dopaminergic basis for well-known behavioral differences in reward and punishment processing between women and men. This has important implications in psychiatric conditions showing sex-specific prevalence rates, altered reward processing and dopamine signaling. The high temporal resolution and pronounced magnitude of task-specific changes make fPET a promising tool to investigate functional neurotransmitter dynamics during cognitive or emotional processing in various brain disorders.

## INTRODUCTION

The processing of reward and punishment represents an essential aspect of one’s mental health. This is reflected in alterations of the reward system in several psychiatric disorders such as addiction, gambling, eating disorders and depression. The prevailing approach to investigate the neural representation of behavioral effects is functional magnetic resonance imaging (fMRI) with the monetary incentive delay (MID) task being the most widely employed paradigm to study reward and punishment processing [1]. Probing differences between monetary gain and loss consistently shows activation of the nucleus accumbens (NAcc) [1], which is the major part of the ventral striatum [2] and pivotal for reward processing [3, 4]. However, blood oxygen level dependent (BOLD) fMRI is directly related to hemodynamic factors and mostly reflects post-synaptic glutamate-mediated signaling [5] instead of mapping specific modulatory neurotransmitter action [6].

Dopamine plays a crucial role in the processing of reward and punishment by specifically encoding these conditions. Animal research has demonstrated that the behavioral response to rewarding and aversive stimuli [7] is mediated by different neuronal projections from the ventral tegmental area to the NAcc [8, 9]. This anatomical separation also implies distinct dopamine signaling that underpin the two motivational signals. In humans endogenous dopamine release can only be assessed indirectly by specific positron emission tomography (PET) radioligands, which compete with the endogenous neurotransmitter to bind at a target receptor. Although the competition model represents a thoroughly validated approach it includes two major disadvantages when investigating human behavior. First, cognitive tasks only yield low signal changes of around 5-15% from baseline [10], even for a recently introduced advancement that offers high temporal resolution [11]. Second, high specificity of observed task effects implies comparison against a control condition, but this in turn requires separate measurements. As a consequence, among those studies investigating dopamine release during monetary gain [12–15] only one also evaluated loss, but without observing significant differences between the two conditions [16].

An important aspect in the context of reward processing and dopamine neurotransmission is the widely described sex difference thereof. Numerous different testing schemes have shown that women are more sensitive to threats and punishment, thus aiming for risk minimization and harm avoidance. However, men tend to opt for greater rewards in terms of money, status and competitive success irrespective of the associated risks [17, 18]. Furthermore, several studies have reported general sex differences of the dopamine system, including ventral tegmental area functioning [19], dopamine synthesis rates at baseline [20] and amphetamine-induced release [21, 22], whereas the latter may only be present in young adults [23]. Nevertheless, differences in reward-specific dopamine release between women and men have not yet been investigated, which is potentially attributable to the methodological difficulties mentioned above. Consequently, the neuronal underpinnings of behavioral sex differences in reward and punishment processing remain largely unknown, particularly because fMRI studies of the MID [1, 24] or other reward paradigms [25, 26] were unable to show any sex differences during reward consumption.

Therefore, the primary aim of this work was to introduce a novel approach, which enables the assessment of rapid changes in dopamine signaling during cognitive performance by extending the technique of functional PET (fPET) imaging [27, 28] to a neurotransmitter level. Here, task-induced functional dynamics of dopamine synthesis were used as an index of dopamine neurotransmission, focusing on the NAcc due to its pivotal role in the processing of reward and punishment. The second aim was to demonstrate the feasibility of this technique by investigating sex differences in the processing of monetary gain and loss on a multimodal level. Thus, we combined task-induced changes in dopamine synthesis with BOLD-derived neuronal activation and modeling of behavioral data to identify the neuronal processes underlying the different behavioral sensitivity to reward and punishment in men and women.

## THEORY

### Synthesis model

To assess task-relevant changes in dopamine signaling during cognitive performance we developed a novel approach, based on the dynamic regulation of neurotransmitter synthesis. As most neurotransmitters cannot pass the blood brain barrier, they are synthetized in the brain through precursor molecules. For dopamine, the main pathway is the conversion of L-tyrosine to L-3,4-dihydroxyphenylalanine (DOPA) via the enzyme tyrosine hydroxylase, and then to dopamine by aromatic amino acid decarboxylase (AADC). Importantly, these enzymes are subject to fast-acting regulatory mechanisms. Tyrosine hydroxylase and AADC activities increase with neuronal firing in order to refill the synaptic vesicles with *de novo* synthetized neurotransmitter after stimulus-induced dopamine release and are further regulated by activation or blockade of dopamine receptors [29–33]. Moreover, the radioligand 6-[^18^F]FDOPA can be incorporated into this synthesis chain, as it is a substrate for AADC, rapidly forming 6-[^18^F]F-dopamine. The radioligand is thus specific to the dopaminergic pathway [34, 35] and represents an established approximation for dopamine synthesis rates [33, 36]. Taken together, the evidence suggests that stimulus-induced activation of dopamine synthesis is also reflected in a proportionally increased radioligand binding (Fig. 1a-b).

**Figure 1:**
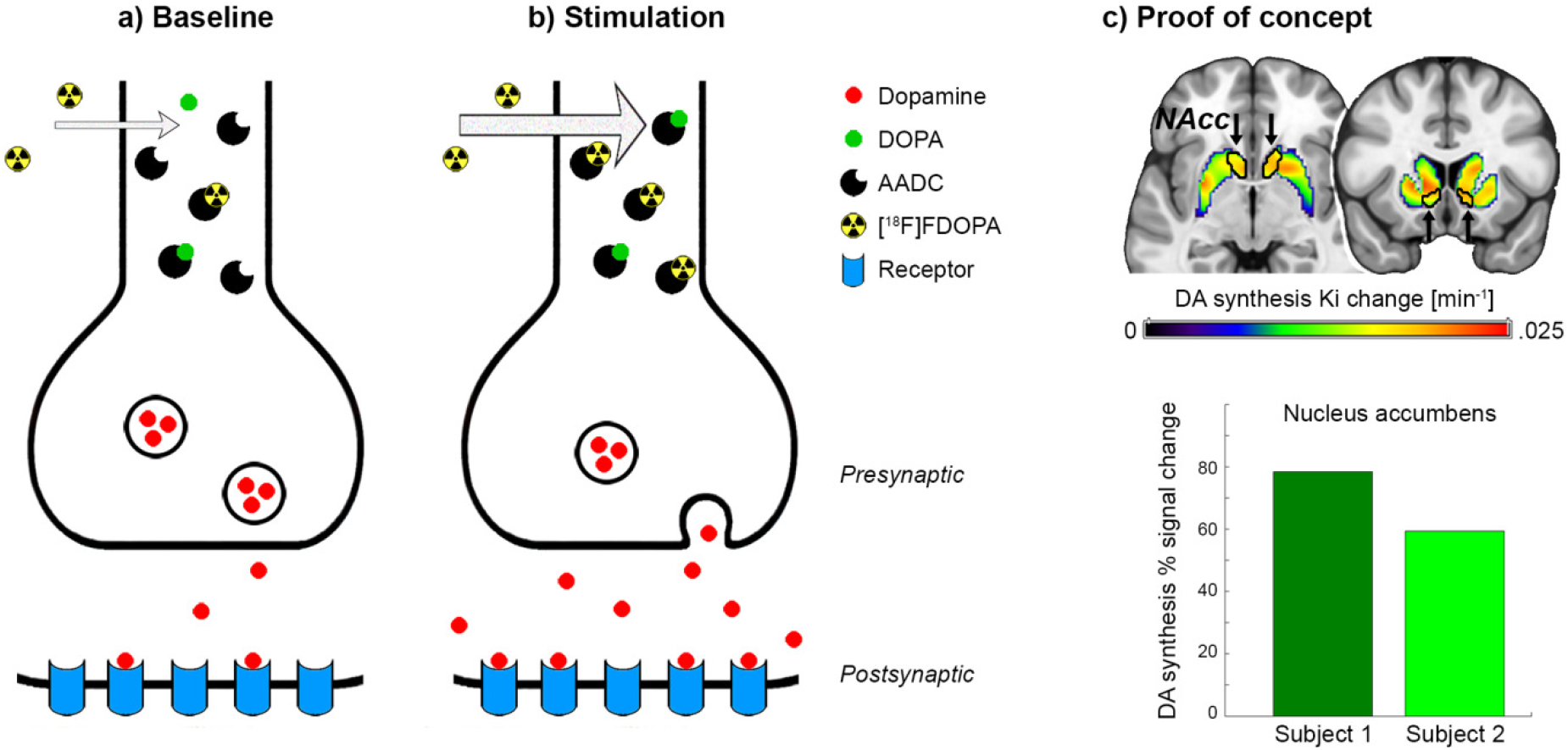
Synthesis model. **a)** The neurotransmitter dopamine (DA) is synthetized from its precursor dihydroxyphenylalanine (DOPA) by the enzyme aromatic amino acid decarboxylase (AADC). Use of the radioligand 6-[^18^F]FDOPA as substrate for AADC is a well-established approach to estimate dopamine synthesis rates at baseline. **b)** Neuronal stimulation leads to dopamine release, but also increases AADC activity to refill synaptic vesicles with *de novo* synthetized neurotransmitter, which in turn is reflected in higher radioligand uptake as indicated by arrow thickness. **c)** The proof of concept experiment showed a marked increase in striatal dopamine synthesis rates Ki during performance of the monetary incentive delay task. The nucleus accumbens (NAcc) region of interest is outlined in black and indicated by arrows, exhibiting 78.4% and 59.4% signal change from baseline for two subjects.

This hypothesis can be directly tested by the application of 6-[^18^F]FDOPA within the framework of functional PET imaging [27, 37]. Similar to fMRI, fPET employs cognitive paradigms in repeated periods of task performance with an alternating control condition, thereby enabling the assessment of task-induced changes of multiple conditions within a single measurement. The radioligand 6-[^18^F]FDOPA is particularly suited for this application, as a bolus + infusion protocol [28] further emphasizes its apparently irreversible binding characteristics [36, 38] (Suppl. Fig. S1b and e), which in turn allows to identify task-specific changes in dopamine synthesis with high temporal resolution.

## MATERIALS and METHODS

### Participants

In total, 41 healthy participants were recruited for this study. Three subjects were excluded as the fPET measurement failed for technical reasons or urinary urgency. Two subjects participated in the proof of concept experiment (age 19.8 and 20.8 years, both female). The main study included 36 participants, who underwent behavioral testing with the MID task (24.5±4.3 years, 18 female). Of those, 16 participants also completed fPET and fMRI examinations (24.8±4.8 years, 7 female). Men and women did not differ regarding their age in the full sample (n=36, p=0.69) or the imaging subsample (n=16, p=0.51). Please see supplement for further details.

After detailed explanation of the study protocol, all participants gave written informed consent. Participants were insured and reimbursed for their participation. The study was approved by the Ethics Committee (ethics number: 2259/2017) of the Medical University of Vienna and procedures were carried out in accordance with the Declaration of Helsinki.

### Cognitive task

Reward and punishment processing was assessed using the well-established [1] and previously employed [39] MID task. Here, participants aim to maximize gain and avoid loss by fast reaction upon presentation of a target stimulus.

As the crucial aspect of the paradigm is the time limit of the reaction, we employed an adaptive algorithm to control the probability for gain and loss. First, the initial reaction time was individually determined directly before each testing procedure (imaging/behavior). Second, the time limit was decreased (increased) during the paradigm if the reaction was fast enough (too slow), to maintain a probability of approximately 0.5. Third, for the main study the time limit was increased (decreased) in the beginning and middle of each task block, which enabled separation of gain and loss by increasing (decreasing) the probability for each condition. The last step allowed assessment of both conditions in a single scan. Please see supplement for a detailed task description.

### Magnetic resonance imaging (MRI)

MRI data was obtained on a 3 T Magnetom Prisma scanner (Siemens Healthineers) using a 64 channel head coil. A structural MRI was acquired with a T1-weighted MPRAGE sequence (TE/TR=2.29/2300 ms, voxel size=0.94 mm isotropic, 5.3 min), which was used to exclude gross neurological abnormalities and for spatial normalization of fPET data. fMRI data was acquired using an EPI sequence (TE/TR = 30/2050 ms, voxel size=2.1 x 2.1 x 2.8 mm + 0.7 mm slice gap).

### Positron emission tomography (PET)

The radioligand was freshly prepared every day by Iason GmbH or BSM Diagnostica GmbH. One hour before start of the fPET measurement, each participant received 150 mg carbidopa p.o. to block peripheral metabolism of the radioligand by aromatic amino acid decarboxylase [36]. fPET imaging was carried out using an Advance PET scanner (GE Healthcare) similar to previously described procedures [27, 28] (see supplement). During the scan the MID task was carried out at 10 (except for the PoC experiments), 20, 30 and 40 min after start of the radioligand application, each lasting for 5 min. Otherwise, a crosshair was presented and subjects were instructed to keep their eyes open and avoid focusing on anything specific (in particular not the task).

### Blood sampling

Arterial blood samples were drawn from the radial artery (see supplement). Manual samples of plasma to whole blood ratio were fitted with a linear function. Correction for radioactive metabolites was based on previous literature, assuming that the only relevant metabolite is 3-O-methyl-6-[^18^F]FDOPA (3-OMFD) after carbidopa pretreatment [36, 38, 40] (see supplement). The final arterial input function was then obtained by multiplication of the whole blood curve with the plasma to whole blood ratio and the parent fraction.

### Quantification of dopamine synthesis rates

Image preprocessing was done as described previously [28] using SPM12 and default parameters unless specified otherwise. fPET images were corrected for head motion (quality=1, registered to mean) and the resulting mean image was coregistered to the T1-weighted structural MRI. The structural scan was spatially normalized to MNI space and the resulting transformation matrices (coregistration and normalization) were applied to the dynamic fPET data. Images were smoothed with an 8 mm Gaussian kernel, masked to include only gray matter voxels and a low-pass filter was applied with the cutoff frequency set to 2.5 min.

The general linear model was used to separate task effects from baseline synthesis (Suppl. Fig. S1c). This included one regressor for each task block (except for the PoC experiments where a single task regressor was used) with a slope of 1 kBq/frame, one representing baseline dopamine synthesis and one for head motion (first principal component of the six motion regressors). The baseline was defined as average time course of all gray matter voxels, excluding those activated in the corresponding fMRI acquisition (contrast success > failure, p<0.001 uncorrected) and those identified in a recent meta-analysis of the MID task (contrasts reward/loss anticipation and reward outcome) [1]. The Gjedde-Patlak plot was then applied to compute the net influx constant Ki as index of dopamine synthesis for baseline and task effects separately (Suppl. Fig. S1e). Task-specific percent signal change (PSC) from baseline was calculated as

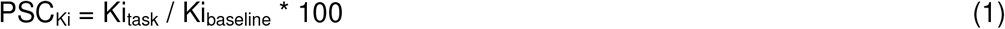

The four task blocks were finally weighted according to task performance (actual gain/possible gain, similar for loss) and averaged to obtain task specific PSC_Ki_ for gain and loss. To assess the specificity of the findings, task-specific changes in dopamine synthesis rates were also calculated independently of baseline Ki values (i.e., not PSC) and without weighting by task performance.

### Neuronal activation

Task-induced neuronal activation was computed as described previously using SPM12 [39]. fMRI BOLD images were corrected for slice timing differences (reference: middle slice) and head motion (quality=1, registered to mean), spatially normalized to MNI space and smoothed with an 8 mm Gaussian kernel. Neuronal activation was estimated across the two runs with the general linear model including one regressor for each cue (gain, loss, neutral), one for the target stimulus and one for each of the potential outcomes (gain, omitted gain, loss, avoided loss, neutral) as well as several nuisance regressors (motion, white matter, cerebrospinal fluid). To obtain an index of reward outcome [1] which is as similar to fPET as possible, parameter estimates were combined as (gain + avoided loss) – (omitted gain + loss). Percent signal changes were computed as

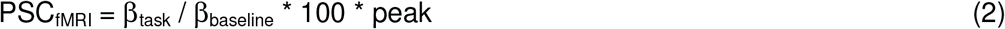

with β_baseline_ and peak representing the constant and the peak value of the fMRI design matrix, respectively [41].

### Statistical analysis

All statistical tests were two-sided and corrected for multiple comparisons with the Bonferroni-Holm procedure (e.g., when testing multiple conditions and/or groups) and the reported p-values have been adjusted accordingly.

For behavioral data, the accumulated amount of money that was gained and lost during the corresponding task blocks of the MID were assessed with one sample t-tests against zero, whereas sex differences were computed by independent samples t-tests. Due to the adaptive nature of the MID task the reaction times were normalized to the mean by subtracting the average reaction time within each block. Differences in reaction times were evaluated by repeated measures ANOVA with the factors sex and amount. Post-hoc t-tests were used to assess sex differences for each amount of money. Furthermore, we modeled the relationship between reaction time and amount with a stepwise linear regression up to 2^nd^ order polynomial functions. Stepwise regression choses the model that best explains the data based on statistical significance. This was done across the entire group (n=36) to test for a general relationship. Subsequently, parameters of the resulting models were also estimated individually for the fPET subjects (n=16) to assess the correlation with task-specific changes in dopamine synthesis using Spearman’s correlation (since n<10 in each group for fPET).

For imaging parameters, the primary region of interest was the NAcc due to its pivotal importance in reward processing [3, 4]. Therefore, values of PSC_Ki_ and PSC_fMRI_ were extracted for this region using the Harvard Oxford atlas as provided in FSL. For comparison, a functional definition of the NAcc was also employed (neuronal activation of reward outcome [1] within the striatum), which comprised 2.55 cm^3^ (in contrast to the NAcc of the Harvard-Oxford atlas with only 1.38 cm^3^). Task-specific changes in dopamine synthesis rates were evaluated by one sample t-tests against zero for gain and loss separately. Similarly, for PSC_Ki_ and PSC_fMRI_ the difference of gain vs. loss was calculated and assessed by one sample t-tests against zero (i.e., being identical to a paired samples t-tests). Finally, sex differences in PSC_Ki_ and PSC_fMRI_ were addressed using an independent samples t-test.

## RESULTS

### Proof of concept (PoC)

To assess the feasibility of the proposed synthesis model an initial PoC experiment was conducted. Two subjects underwent fPET imaging with the radioligand 6-[^18^F]FDOPA while performing the MID task. In both subjects the task induced substantial changes in NAcc dopamine synthesis of 78.4% and 59.4% from baseline (Fig. 1c), supporting the feasibility of the approach to assess task-specific changes in dopamine neurotransmission.

### Behavioral data

Since the PoC experiment combined monetary gain and loss within a task block, the main study specifically aimed to disentangle these two effects on a behavioral (n=36) and neurobiological level (n=16, see below). The task was extended to 4 blocks and each of them manipulated to enable the separate assessment of monetary gain and loss.

Behavioral data showed that average monetary gain and loss were significantly different from zero (all t=10.0 to 12.6, p=1.7*10^−8^ to 2*10^−9^, Fig. 2a), indicating successful task manipulation. Women gained significantly more than men (5.6±2.4 € vs. 4.2±1.7 €, t=2.1, p=0.047), but both groups showed similar loss (−5.3±1.9 € vs. −5.5±1.9 €, p=0.8).

**Figure 2:**
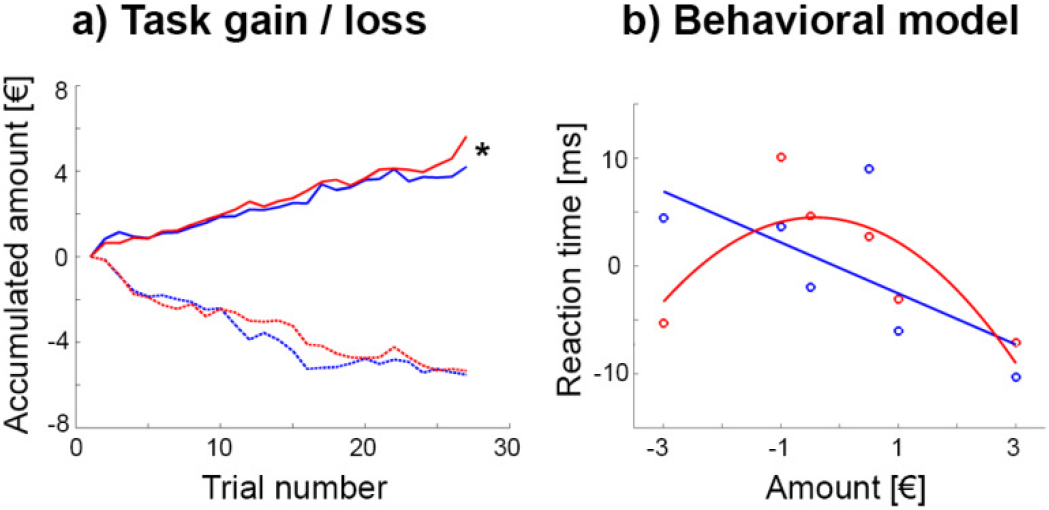
Behavioral data (blue = men, red = women). **a)** The monetary incentive delay task was manipulated by modifying the reaction time limit for a successful trial completion, which enabled separate assessment of monetary gain and loss. During the gain task block women earned significantly more money than men (5.6±2.4 € vs. 4.2±1.7 €, t=2.1, *p=0.047), but both groups showed similar loss (−5.3±1.9 € vs. −5.5±1.9 €, p=0.8). **b)** The association between the individually normalized reaction times and the gained/lost amount of money was modelled by a linear relationship in men (reaction time = −0.18 - 2.36*amount, p_linear_=0.0012, p_quadratic_=0.11), where a steeper negative slope indicated faster reaction time and thus higher sensitivity for reward. In contrast, the association was characterized by an inverted u-shaped function in women (reaction time = 4.31 - 0.95*amount - 1.16*amount^2^, p_linear_=0.2, p_quadratic_=0.001). Since these two functions exhibit the most pronounced difference for high amounts of loss, a strong negative quadratic term was interpreted as high sensitivity for punishment. Circles denote average values for each amount and lines are model fits across the entire data set (n=36 subjects), reaction times are mean centered due to the adaptive nature of the MID task.

The difference in monetary gain was also reflected in the normalized reaction times, with a main effect of sex (F_(1,34)_=6.9, p=0.013) and amount (F_(5,170)_=4.4, p=0.0009) as well as a trend for an interaction effect sex * amount (F_(5,170)_=2.0, p=0.08). Post-hoc t-test indicated that this seemed to be driven by the −3 € condition with women showing a faster reaction than men (t=2.0, p=0.049).

We further aimed to model the behavioral response in more detail, as the relationship between reaction time and amount for each group. In men, this was best described by a negative linear function (reaction time = −0.18 - 2.36*amount, p_linear_=0.0012, p_quadratic_=0.11), with a faster reaction for higher monetary gains (Fig. 2b). In contrast, the association in women was characterized by an inverted u-shaped function (reaction time = 4.31 - 0.95*amount - 1.16*amount^2^, p_linear_=0.2, p_quadratic_=0.001), with faster reaction times for high amounts of loss as compared to men. Thus, we interpreted the linear (quadratic) term for men (women) as index for reward (punishment) sensitivity, i.e., the more negative the parameter, the faster the reaction time for high gain (loss).

### Functional dynamics in dopamine synthesis

To assess reward-specific changes in dopamine synthesis, 16 of the above subjects also underwent fPET with the radioligand 6-[^18^F]FDOPA (7 female). The MID task yielded increased dopamine synthesis rates in the NAcc during gain (men: 77.6±32.9% from baseline, women: 51.2±16.5%) and loss (men: 49.4±26.7%, women: 78.4±18.6, all t=5.5 to 11.1, all p=0.0001 to 0.0005, Fig. 3b). As a result, the direct comparison between the two conditions showed higher dopamine synthesis rates in men for gain vs. loss (28.2±42.6%, t=2.0, p=0.08). Interestingly, the direction of this difference was reversed in women with higher dopamine synthesis during loss vs. gain (−27.3±17.2%, t=−4.2, p=0.01, Fig. 4a).

**Figure 3:**
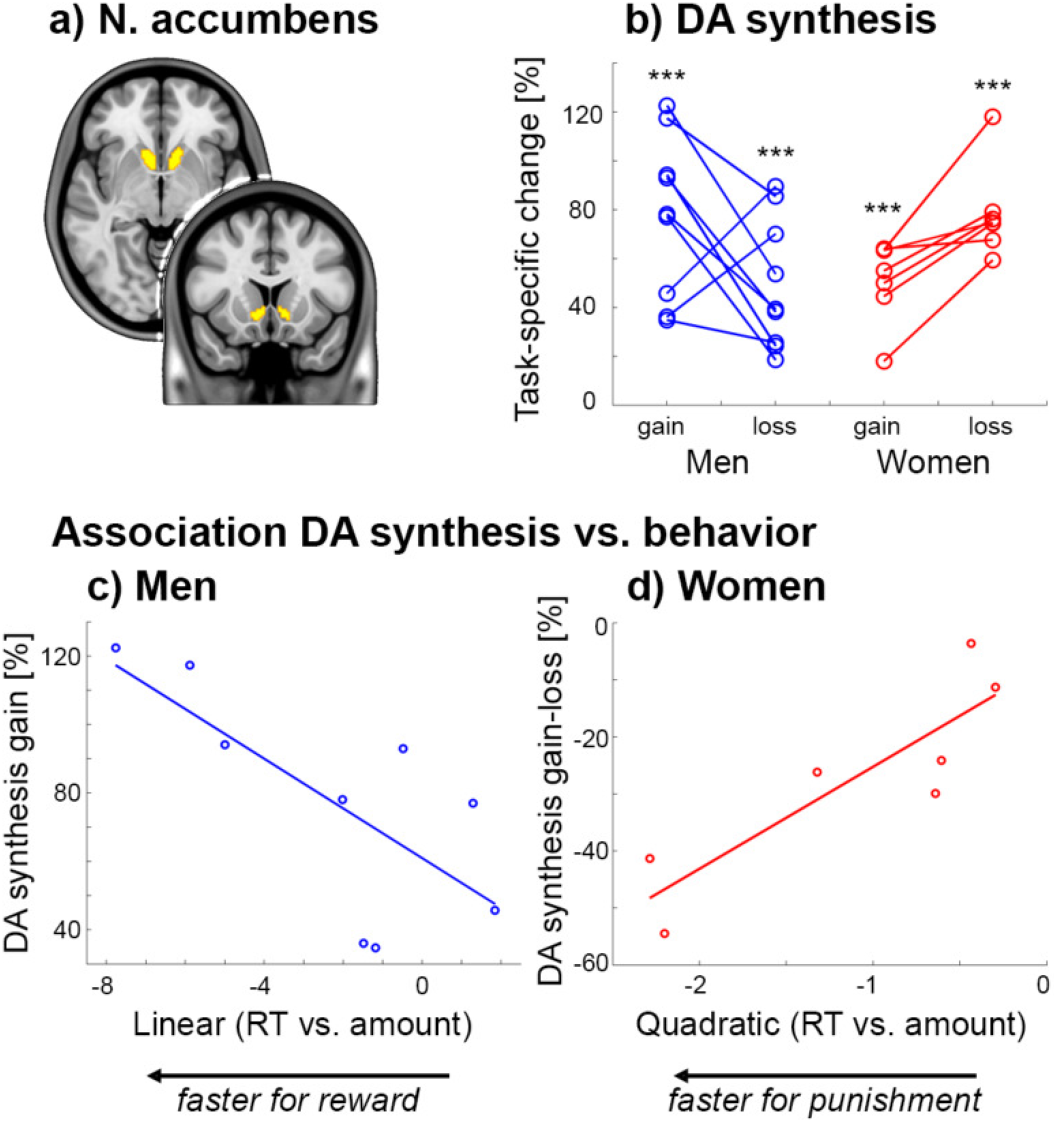
Functional PET imaging of task-specific dopamine synthesis. **a)** Region of interest of the nucleus accumbens (NAcc) from the Harvard-Oxford atlas. **b)** Processing of monetary gain and loss resulted in pronounced changes in NAcc dopamine synthesis (***all p<0.001). While men showed higher dopamine synthesis changes for gain vs. loss (n = 9), women exhibited the opposite pattern (n=7, see also figure 4a). **c-d)** The individually modelled associations between reaction time (RT) and amount were used as indices for reward and punishment sensitivity in men (linear term) and women (quadratic term), respectively (see figure 2b). These behavioral indices showed an association with task-specific changes in NAcc dopamine synthesis during monetary gain in men (c) rho=−0.7, p=0.043) and the difference between gain and loss in women (d) rho=0.89, p=0.012).

**Figure 4:**
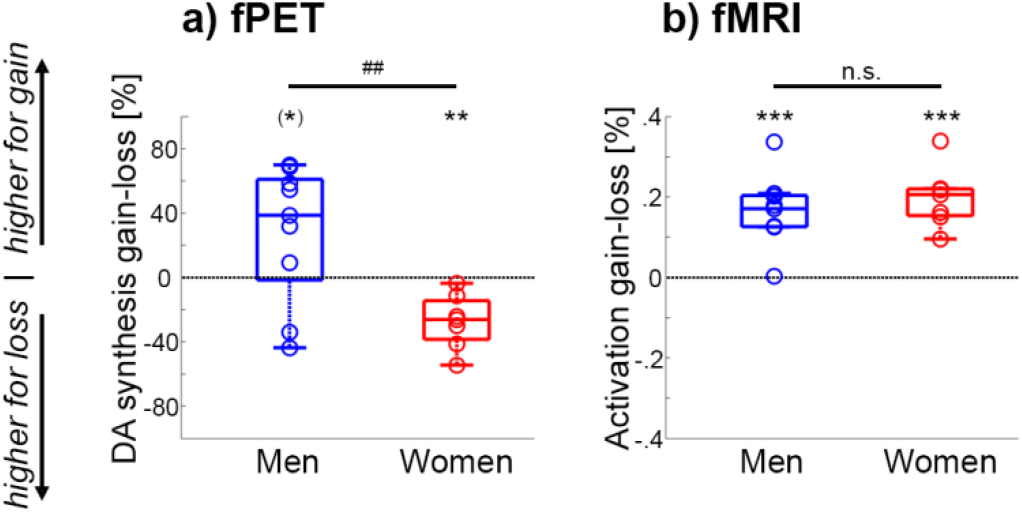
Comparison between fPET and fMRI. **a)** In men the task-specific changes in NAcc dopamine synthesis were higher for gain than for loss (^(*)^p=0.08). In contrast, women showed the opposite pattern with higher changes in dopamine synthesis during loss vs. gain (**p<0.01), leading to a significant difference between the two groups (t=3.2, ^##^p=0.006). **b)** Although neuronal activation obtained with BOLD fMRI indeed showed robust NAcc signal changes for the contrast gain vs. loss for men and women (t=5.5 to 6.9, ***p=0.0005 to 0.0006), there was no significant difference between the two groups (p=0.4). Boxplots indicate median values (center line), upper and lower quartiles (box limits) and 1.5*interquartile range (whiskers).

Proceeding from the distinct models to characterize the behavioral response of monetary gain and loss (Fig. 2b), we assessed the relationship between individual model parameters (linear and quadratic terms) and task-specific dopamine synthesis rates. This resulted in a significant association in men between the linear term and NAcc dopamine synthesis during gain (n=9, rho=−0.7, p=0.043, Fig. 3c). On the other hand, the quadratic term in women was positively associated with NAcc dopamine synthesis of gain vs. loss (n=7, rho=0.89, p=0.012, Fig. 3d).

### Sex differences

Finally, NAcc dopamine synthesis rates between gain vs. loss were significantly higher for men than women (t=3.2, p=0.006, Fig. 4a). This sex difference was similarly present when using a functional delineation of the NAcc [1] (t=2.7, p=0.02), for raw dopamine synthesis values (i.e., not % signal change from baseline, t=3.3, p=0.005) and without weighting by task performance (t=3.0, p=0.01). Furthermore, the sex difference was specific for the NAcc as exploratory analysis revealed similar dopamine synthesis rates for men and women in the caudate (p=0.8) and putamen (p=0.2).

For direct comparison we also assessed neuronal activation, where the same subjects as in the fPET experiment also underwent fMRI. In line with previous reports [1] we observed robust neuronal activation in the NAcc for gain vs. loss in men and women (t=5.5 to 6.9, p=0.0005 to 0.0006, Fig. 4b). As expected, there was however no significant sex difference in activation for the atlas-based (p=0.4) or the functional delineation of the NAcc (p=0.3).

Similar to a previous study [20], we also observed a sex difference in *baseline* NAcc dopamine synthesis (men: Ki=0.017±0.002 min^−1^, women: Ki=0.02±0.002 min^−1^, t=4.4, p=0.0007). It is however unlikely that these baseline differences affect the task-specific estimates (see limitations).

## DISCUSSION

In this work we introduce a novel framework for the assessment of task-specific changes in dopamine neurotransmission, which is based on the dynamic regulation of neurotransmitter synthesis quantified by functional PET imaging. Processing of monetary gain and loss induced robust changes in dopamine signaling in the living human brain even for the direct comparison of these two conditions, demonstrating the high sensitivity and specificity of the approach. Crucially, task-induced changes in dopamine synthesis showed sex-specific differences in the opposite direction with higher synthesis rates in men for gain vs. loss but vice versa in women, directly reflecting behavioral sex differences in reward and punishment sensitivity. Since this sex difference was not present in common BOLD-derived assessment of neuronal activation, our findings have important implications for the interpretation of numerous fMRI studies on reward processing. This is also essential in various clinical populations, where the sex-specific influence on the link between altered reward processing and dopamine signaling is not yet fully understood [42].

The current work provides a biological basis for the well-known behavioral differences in reward and punishment sensitivity between men and women [17, 18]. We hereby extend general sex differences of the dopamine system [20–22] specifically to the processing of gain and loss and directly link changes in dopamine neurotransmission with the corresponding behavioral response. This is also supported by pharmacological effects observed in animals and humans. For instance, male rats aim for large rewards independent of the risk, whereas females decrease such choices in order to avoid punishment. This sex difference was even more pronounced by the dopamine releasing agent amphetamine, where females abolished the choice for risky rewards to a much larger extent than males [43]. On the other hand, studies in humans have shown that men often opt for selfish rewards, but women take more prosocial choices. However, pharmacological blockade of dopamine D2/D3 receptors shifts these preferences and thereby eliminate the sex difference in prosocial choices, i.e., men and women showed similar preference for selfish rewards [44]. Taken together, these findings suggest that sex differences in reward behavior are substantially driven by dopamine neurotransmission. It is worth to note that pharmacological challenges may represent an unspecific assessment of neurotransmitter action. The systemic manipulation affects the entire brain, possibly eliciting complex downstream effects, and the use of potent challenge agents may overshadow subtle physiological and behavioral differences. Therefore, our results provide novel evidence in this context through the direct and spatially targeted assessment of reward-specific dopamine signaling itself, without manipulation of the neurotransmitter system. This enabled us to disentangle the dopaminergic involvement in monetary gain and loss, which revealed opposing changes in synthesis rates between men and women.

In contrast, such an evaluation was not accessible by previous approaches (see introduction for PET findings on the competition model), including reward-specific neuronal activation obtained with fMRI. Again, it needs to be emphasized that neither this nor previous fMRI studies [1, 24, 25] revealed any (and particularly not opposing) sex differences in NAcc activation between gain and loss. fMRI based on the BOLD signal is dependent on the link between neuronal activation and changes in hemodynamic factors such as blood flow, volume and oxygenation [6, 45]. Blood flow is locally controlled by the major neurotransmitter glutamate, and thus it is widely accepted that the BOLD signal mostly reflects postsynaptic glutamate-mediated signaling [5, 46]. Although monoamine neurotransmitters such as dopamine may also modulate blood flow [47], this does not seem to translate into corresponding fMRI signal changes, at least for the processing of monetary gain and loss using the widely employed MID task. We acknowledge that previous work has indicated a relationship between dopamine release and fMRI [48, 49], but these were again based on potent pharmacological manipulations, which may not be directly comparable to more subtle cognitive effects (see above). Instead, it appears that during cognitive task performance the limited contribution of dopamine to the BOLD signal gets lost in major downstream effects of glutamate action [5] that regulate blood flow. We speculate that the latter two are not sufficiently specific [6] to identify sex differences in neuronal activation during reward processing. This may have substantial implications for the investigation of several brain disorders with dopamine dysfunction such as addiction, schizophrenia or depression, where fMRI represents one of the most widely used methods. Our results suggest that BOLD signal alterations may not primarily reflect the underlying dopaminergic changes, especially when investigating the reward system in men and women. Further work is required to elucidate the exact difference that cognitive and pharmacological stimulation exert on the relationship between BOLD imaging and dopamine signaling and if this extends beyond sex differences of reward processing.

Although not directly assessed, there are two essential lines of evidence which strongly support the concept that task-specific changes in the 6-[^18^F]FDOPA signal are related to dopamine release. As mentioned, dopamine synthesis is subject to fast-acting regulatory mechanisms, which is activated by neuronal firing to refill the synaptic vesicles [29–31]. Moreover, dopamine synthesis is also increased by the dopamine releasing agent amphetamine as demonstrated in rats [50] and monkeys using PET [51]. Notably, a previous study reported no relationship between dopamine synthesis and release [52], but it is important to mention that synthesis was only investigated at baseline (i.e., without any task- or drug-induced stimulation). In contrast, we specifically assessed changes in dopamine synthesis during task performance and thus the previous finding is not in contrast to the synthesis model. Hence, the herein proposed approach offers an alternative to the competition model as the crucial factor to identify task-specific changes is the incorporation of radioligands into the dynamic regulation of enzymes responsible for neurotransmitter synthesis (instead of direct competition between radioligand and endogenous neurotransmitter).

The different neurobiological basis of these two approaches (i.e., competition vs. synthesis model) seems to explain the marked signal changes observed during the reward task of around 75% from baseline and 25% for the direct comparison of gain vs. loss. This underlines the high sensitivity of the technique but also the high specificity, with the ability to separate subtle effects of behaviorally similar conditions. Furthermore, fPET allows to assess task-specific changes of multiple conditions in a single within-scan design, thereby eliminating intrasubject variability related to differences in habituation, motivation or performance of repeated measurements. These advantages seem to translate into robust effects even with a low sample size, thereby mitigating the limitation of the current study that imaging was only performed in a subset of the cohort. Another limitation is the use of a literature-based correction for radioactive metabolites instead of an individual one. Although this may indeed change the absolute values of dopamine synthesis to a certain extent, it does not influence the reward-specific effects. Again, in a within-scan design any “global” parameter will affect baseline and task-specific synthesis rates in an equal manner and will thus cancel out when calculating percent signal change or differences between gain and loss. This applies for instance to radioactive metabolites as well as sex differences in dopamine synthesis at baseline [20]. Regarding the latter issue, baseline differences (if at all) would most likely cause general differences in task-specific dopamine synthesis across all task conditions. However, the observed task-specific changes were higher in men than women for gain, but vice versa for loss, which argues against a dependency of task estimates on baseline synthesis.

To summarize, the current work provides a strong motivation for further investigations of functional neurotransmitter dynamics during cognitive processing. The framework of fPET imaging offers important advantages of high temporal resolution, robust effect size of task-induced changes and the possibility to assess multiple task conditions in a single measurement. Future studies should aim for an in-depth evaluation of stimulus-dependent activation of dopamine synthesis, proceeding from previous findings which link neurotransmitter synthesis and release [50, 51]. Moreover, our results suggest that reward-specific neuronal activation should not unequivocally be interpreted as corresponding changes in dopamine signaling and that the investigation of sex differences in this context requires further attention. This may be of pivotal relevance for the assessment of psychiatric populations such as addictive and gambling disorders or depression, given the different prevalence rates in men and women as well as alterations in reward processing and dopamine signaling. The introduced approach enables to address important future questions of human cognition and to investigate whether the observed reward- and sex-specific differences in dopamine synthesis will translate to clinically relevant characteristics for patient diagnosis or treatment.

## ETHICS APPROVAL

The study was approved by the Ethics Committee (ethics number: 2259/2017) of the Medical University of Vienna and procedures were carried out in accordance with the Declaration of Helsinki.

## CONSENT TO PARTICIPATE

After detailed explanation of the study protocol, all participants gave written informed consent. Participants were insured and reimbursed for their participation.

## CONSENT FOR PUBLICATION

Not applicable.

## AVAILABILITY OF DATA, MATERIAL AND CODE

Raw data cannot be shared due to data protection laws. Processed data and code are available from the corresponding author upon reasonable request.

## AUTHOR CONTRIBUTIONS

Study design: A.H., W.W., M.H., R.L.

Data acquisition: M.B.R., L.R., A.H., V.P., P.M., G.M.G.

Data analysis: A.H., M.B.R., L.R.

Manuscript preparation: A.H., L.R., M.B.R., R.L.

All authors discussed the implications of the findings and approved the final version of the manuscript.

## ACKNOWLEDGEMENTS

The authors are particularly grateful to S. Oldham and V. Lorenzetti for providing the meta-analysis maps of the monetary incentive delay task. We would like to thank P. Baldinger-Melich and A. Basaran for medical and measurement support as well as V. Ritter and the medical students of the Neuroimaging Labs for subject recruitment and administrative support.

## SUPPLEMENTARY MATERIAL

### MATERIALS and METHODS

#### Participants

General health of all participants was ensured at the screening visit by an experienced psychiatrist as based on the subjects’ medical history. Exclusion criteria were current and previous (12 months) somatic, neurological or psychiatric disorders, current and previous substance abuse or psychotropic medication. For participants who underwent fPET and fMRI a more extensive medical examination was carried out, including blood tests, electrocardiography, neurological testing and the Structural Clinical Interview for DSM-IV. Additional exclusion criteria were contraindications for MRI scanning, previous study-related radiation exposure (10 years), pregnancy or breast feeding. Thus, female participants underwent a urine pregnancy test at the screening visit and before the fPET and MRI scans.

#### Cognitive task

The task was designed in an even-related manner. Each trial started with the presentation of the potential gain or loss (e.g., +3 €, −1 €) for an unknown variable duration of 3 - 5 s (0.5 s steps, uniformly distributed). After that the target stimulus (!) was shown and subjects were required to press a button as fast as possible. If the reaction was within a given time limit the amount was gained or loss was avoided. Otherwise, the amount was not gained or lost. Each button press was followed by immediate feedback (2 s), showing the amount gained or lost, the outcome of the reaction (in green for success, red for failure) and the current account sum. Trials were separated by a crosshair with a variable duration of 3 - 7 s (1 s steps, uniformly distributed). To maintain a high level of attention and to enable modeling of the behavioral response, 6 different amounts of money were used for the task (0.5, 1 and 3 € each for gain and loss, initial amount was 10 €). Furthermore, motivation of the participants was kept high by the instruction that the final amount was paid out in addition to a fixed reimbursement.

The crucial aspect of this paradigm is the time limit of the reaction for the button press, where we employed an adaptive algorithm. For fPET and behavioral testing the task was carried out in several blocks, thus, the initial time limit of each block was set to the median of the reaction times of the previous block. The time limit for the first block (and for fMRI) was determined as median of 8 trials taken right before the start of the experiment in the scanner. Second, the time limit for each of the different amounts was adaptively decreased (increased) within a task block by 50 ms if the reaction was fast enough (too slow). These settings maintain an average probability of approximately 0.5 to gain or lose a certain amount.

For the proof of concept (PoC) experiments, 3 task blocks of 5 min were carried out each with 27 trials (equal distribution of amounts presented in random order) without further manipulation of the reaction time limits (i.e., probability of 0.5 for gain and loss, Suppl. Fig. S1a).

For the main study, 4 task blocks of 5 min were carried out in a similar manner, but the probability for monetary gain and loss was manipulated within each block by changing the reaction time limit (Suppl. Fig. S1a). For each of the 2 blocks of monetary gain (loss), the initial time limits of all amounts were increased (decreased) by 50 ms. Furthermore, in the middle of each block the time limits were reset to the median of the preceding reaction times and 25 ms were again added (subtracted). This enabled the separate assessment of gain and loss even within the lower temporal resolution of fPET. The two conditions were presented in alternating order with randomization of the starting condition. Of note, monetary “gain” in this setting combines actual gain with avoided loss, whereas “loss” represents actual loss and omitted gain. The requirement for such a design is however well supported by previous studies indicating that avoided loss represents a relative reward [1]. For the behavioral testing, participants completed the same task version as for fPET but outside the scanner. For the fMRI measurements the MID task was carried out in 2 runs of 8.2 min each.

The manipulation of the time limits (i.e., probability for gain and loss) was similar to that of fPET, but the trial order was randomized to avoid blocks of continuous gain or loss since fMRI has a higher temporal resolution and allows to model each trial separately. Furthermore, a neutral condition was included where no money was at stake (0 €). For all task versions, participants were blind to the adaption and manipulation of the time limits. This was also ensured by the design, where the probability for gain and loss is individually controlled via the time limit, since overruling the participant’s actual reaction time would be easily recognized. The task was implemented in Psychtoolbox v3.0.12.

#### Positron emission tomography (PET)

To minimize head movement each participant’s head was placed in a cushioned polyurethane bowl with straps around the forehead. The paradigm was visualized on a common LCD screen and presented to the subjects by a mirror, which was placed in front of the participant’s eyes using a custom-made wooden construction. Attenuation correction was performed for 5 min with retractable ^68^Ge rod sources, which also included the mirror construction. Dynamic fPET acquisition in 3D-mode started with the intravenous bolus + infusion of the radioligand 6-[^18^F]FDOPA. The injected dose was 5.5 MBq/kg body weight and the measurement time was 50 min. To increase the signal to noise ratio, 20% of the dose was given as bolus for 1 min [2] and the remainder as constant infusion for the rest of the scan (89 kBq/kg/min) using a perfusion pump (Syramed μSP6000, Arcomed, Regensdorf, Switzerland). fPET images were then reconstructed to frames of 43 s, yielding 70 frames in total and 7 frames for each task block.

#### Blood sampling

Automatic sampling was carried out for the first 5 min (4 ml/min, Allogg, Mariefred, Sweden). Manual samples were taken at 3, 4, 5, 16, 26, 36 and 46 min, i.e., at periods of task pauses. For the manually obtained samples, activity in whole blood and plasma (after centrifugation) were measured in a gamma counter (Wizward^2^, 3”, Perkin Elmer). Automatic and manual samples were then combined, were the first manual samples served for a measurement-specific calibration between the automated sampling system and the gamma counter.

The 3-OMFD fraction was extracted from previous studies [3, 4] and fitted with a single exponential function (Suppl. Fig. S1d). Since this represents the metabolite fraction after a bolus application, we adapted the function to the herein employed radioligand application protocol as the sum of an initial 20% bolus and proportionally lower boli administered every further minute (Suppl. Fig. S1d). We are aware that a literature-based metabolite correction may affect the individual estimates of dopamine synthesis rates. However, it equally affects task-specific changes thereof as they are acquired within the same measurement. Thus, the individual variation in metabolism will cancel out when calculating percent signal change from baseline and differences between task conditions (see also discussion). Moreover, the bolus + infusion protocol resulted in a reduction of the 3-OMFD fraction by 51.4% (area under the curve), further minimizing the influence of radioactive metabolites.

## SUPPLEMENTARY FIGURES

**Supplementary figure S1:**
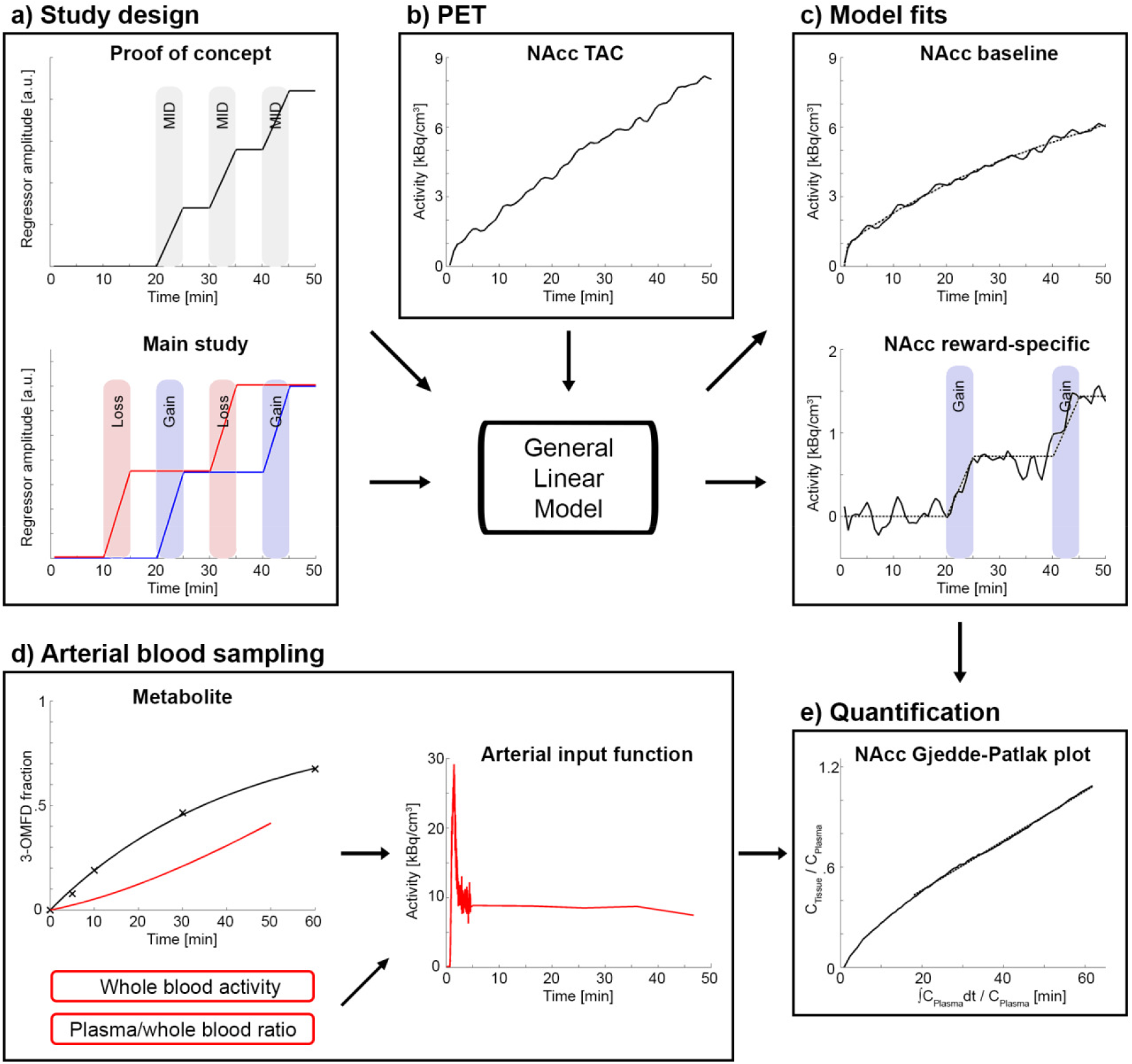
fPET analysis workflow. **a)** During the PET scan the monetary incentive delay (MID) task was carried out in blocks of 5 min. In the proof of concept experiment 3 blocks were employed without a separation of gain and loss conditions and an average probability for success of 0.5. For the main study 4 blocks were completed and the MID task was manipulated to disentangle gain and loss by increasing (decreasing) the probability for success (failure) within the corresponding blocks. **b)** PET measurements were carried out using the radioligand 6-[^18^F]FDOPA. The bolus + infusion protocol emphasized the irreversibly uptake of the radioligand. **c)** Time activity curves (TAC) were then modeled according to the study design to separate baseline and task-specific effects. Model fits (dotted line) indicate a robust increase in radioligand uptake during the task condition in the nucleus accumbens (NAcc). **d)** The metabolite fraction (3-OMFD) was extracted from previous studies [3, 4], fitted with a single exponential function (black line) and adapted to match the bolus + infusion protocol (red line). This was combined with the individually measured whole blood activity and plasma to whole blood ratio to obtain the arterial input function, which reaches steady state approximately after 5 min. **e)** Quantification was carried out with the Gjedde-Patlak plot, yielding dopamine synthesis rates at baseline and for each task condition. All data for the TAC (b), model fits (c), arterial input function (d) and quantification (e) were extracted from a representative subject.

## REFERENCES

1. Oldham S, Murawski C, Fornito A, Youssef G, Yucel M, Lorenzetti V. The anticipation and outcome phases of reward and loss processing: A neuroimaging meta-analysis of the monetary incentive delay task. Hum Brain Mapp. 2018;39:3398–418.

2. Neto LL, Oliveira E, Correia F, Ferreira AG. The human nucleus accumbens: where is it? A stereotactic, anatomical and magnetic resonance imaging study. Neuromodulation. 2008;11:13–22.

3. Berridge KC, Kringelbach ML. Affective neuroscience of pleasure: reward in humans and animals. Psychopharmacology (Berl). 2008;199:457–80.

4. Haber SN, Knutson B. The reward circuit: linking primate anatomy and human imaging. Neuropsychopharmacology. 2010;35:4–26.

5. Attwell D, Laughlin SB. An energy budget for signaling in the grey matter of the brain. J Cereb Blood Flow Metab. 2001;21:1133–45.

6. Heeger DJ, Ress D. What does fMRI tell us about neuronal activity? Nat Rev Neurosci. 2002;3:142–51.

7. Bergamini G, Sigrist H, Ferger B, Singewald N, Seifritz E, Pryce CR. Depletion of nucleus accumbens dopamine leads to impaired reward and aversion processing in mice: Relevance to motivation pathologies. Neuropharmacology. 2016;109:306–19.

8. Matsumoto M, Hikosaka O. Two types of dopamine neuron distinctly convey positive and negative motivational signals. Nature. 2009;459:837–41.

9. Cohen JY, Haesler S, Vong L, Lowell BB, Uchida N. Neuron-type-specific signals for reward and punishment in the ventral tegmental area. Nature. 2012;482:85–8.

10. Egerton A, Mehta MA, Montgomery AJ, Lappin JM, Howes OD, Reeves SJ, et al. The dopaminergic basis of human behaviors: A review of molecular imaging studies. Neurosci Biobehav Rev. 2009;33:1109–32.

11. Lippert RN, Cremer AL, Edwin Thanarajah S, Korn C, Jahans-Price T, Burgeno LM, et al. Time-dependent assessment of stimulus-evoked regional dopamine release. Nat Commun. 2019;10:336.

12. Schott BH, Minuzzi L, Krebs RM, Elmenhorst D, Lang M, Winz OH, et al. Mesolimbic functional magnetic resonance imaging activations during reward anticipation correlate with reward-related ventral striatal dopamine release. J Neurosci. 2008;28:14311–9.

13. Joutsa J, Johansson J, Niemela S, Ollikainen A, Hirvonen MM, Piepponen P, et al. Mesolimbic dopamine release is linked to symptom severity in pathological gambling. Neuroimage. 2012;60:1992–9.

14. Jonasson LS, Axelsson J, Riklund K, Braver TS, Ogren M, Backman L, et al. Dopamine release in nucleus accumbens during rewarded task switching measured by [(1)(1)C]raclopride. Neuroimage. 2014;99:357–64.

15. Weiland BJ, Heitzeg MM, Zald D, Cummiford C, Love T, Zucker RA, et al. Relationship between impulsivity, prefrontal anticipatory activation, and striatal dopamine release during rewarded task performance. Psychiatry Res. 2014;223:244–52.

16. Pappata S, Dehaene S, Poline JB, Gregoire MC, Jobert A, Delforge J, et al. In vivo detection of striatal dopamine release during reward: a PET study with [(11)C]raclopride and a single dynamic scan approach. Neuroimage. 2002;16:1015–27.

17. Cross CP, Copping LT, Campbell A. Sex differences in impulsivity: a meta-analysis. Psychol Bull. 2011;137:97–130.

18. Torrubia R, Avila C, Molto J, Caseras X. The Sensitivity to Punishment and Sensitivity to Reward Questionnaire (SPSRQ) as a measure of Gray’s anxiety and impulsivity dimensions. Pers Indiv Differ. 2001;31:837–62.

19. Gillies GE, Virdee K, McArthur S, Dalley JW. Sex-dependent diversity in ventral tegmental dopaminergic neurons and developmental programing: A molecular, cellular and behavioral analysis. Neuroscience. 2014;282:69–85.

20. Laakso A, Vilkman H, Bergman J, Haaparanta M, Solin O, Syvalahti E, et al. Sex differences in striatal presynaptic dopamine synthesis capacity in healthy subjects. Biol Psychiatry. 2002;52:759–63.

21. Munro CA, McCaul ME, Wong DF, Oswald LM, Zhou Y, Brasic J, et al. Sex differences in striatal dopamine release in healthy adults. Biol Psychiatry. 2006;59:966–74.

22. Riccardi P, Zald D, Li R, Park S, Ansari MS, Dawant B, et al. Sex differences in amphetamine-induced displacement of [(18)F]fallypride in striatal and extrastriatal regions: a PET study. Am J Psychiatry. 2006;163:1639–41.

23. Smith CT, Dang LC, Burgess LL, Perkins SF, San Juan MD, Smith DK, et al. Lack of consistent sex differences in D-amphetamine-induced dopamine release measured with [(18)F]fallypride PET. Psychopharmacology (Berl). 2019;236:581–90.

24. Rademacher L, Krach S, Kohls G, Irmak A, Grunder G, Spreckelmeyer KN. Dissociation of neural networks for anticipation and consumption of monetary and social rewards. Neuroimage. 2010;49:3276–85.

25. Weafer J, Crane NA, Gorka SM, Phan KL, de Wit H. Neural correlates of inhibition and reward are negatively associated. Neuroimage. 2019;196:188–94.

26. van den Bos R, Homberg J, de Visser L. A critical review of sex differences in decision-making tasks: focus on the Iowa Gambling Task. Behav Brain Res. 2013;238:95–108.

27. Hahn A, Gryglewski G, Nics L, Hienert M, Rischka L, Vraka C, et al. Quantification of Task-Specific Glucose Metabolism with Constant Infusion of 18F-FDG. J Nucl Med. 2016;57:1933–40.

28. Rischka L, Gryglewski G, Pfaff S, Vanicek T, Hienert M, Klobl M, et al. Reduced task durations in functional PET imaging with [(18)F]FDG approaching that of functional MRI. Neuroimage. 2018;181:323–30.

29. Morgenroth VH, 3rd, Boadle-Biber M, Roth RH. Tyrosine hydroxylase: activation by nerve stimulation. Proc Natl Acad Sci U S A. 1974;71:4283–7.

30. Neff NH, Hadjiconstantinou M. Aromatic L-amino acid decarboxylase modulation and Parkinson’s disease. Prog Brain Res. 1995;106:91–7.

31. Meiser J, Weindl D, Hiller K. Complexity of dopamine metabolism. Cell Commun Signal. 2013;11:34.

32. Cumming P, Ase A, Laliberte C, Kuwabara H, Gjedde A. In vivo regulation of DOPA decarboxylase by dopamine receptors in rat brain. J Cereb Blood Flow Metab. 1997;17:1254–60.

33. Barrio JR, Huang SC, Phelps ME. Biological imaging and the molecular basis of dopaminergic diseases. Biochem Pharmacol. 1997;54:341–8.

34. Doudet DJ, Chan GLY, Holden JE, McGeer EG, Aigner TA, Wyatt RJ, et al. 6-[F-18]fluoro-L-DOPA PET studies of the turnover of dopamine in MPTP-induced parkinsonism in monkeys. Synapse. 1998;29:225–32.

35. Rinne JO, Nurmi E, Ruottinen HM, Bergman J, Eskola O, Solin O. [F-18]FDOPA and [F-18]CFT are both sensitive PET markers to detect presynaptic dopaminergic hypofunction in early Parkinson’s disease. Synapse. 2001;40:193–200.

36. Cumming P, Gjedde A. Compartmental analysis of dopa decarboxylation in living brain from dynamic positron emission tomograms. Synapse. 1998;29:37–61.

37. Villien M, Wey HY, Mandeville JB, Catana C, Polimeni JR, Sander CY, et al. Dynamic functional imaging of brain glucose utilization using fPET-FDG. Neuroimage. 2014;100:192–9.

38. Holden JE, Doudet D, Endres CJ, Chan GL, Morrison KS, Vingerhoets FJ, et al. Graphical analysis of 6-fluoro-L-dopa trapping: effect of inhibition of catechol-O-methyltransferase. J Nucl Med. 1997;38:1568–74.

39. Pfabigan DM, Seidel EM, Sladky R, Hahn A, Paul K, Grahl A, et al. P300 amplitude variation is related to ventral striatum BOLD response during gain and loss anticipation: an EEG and fMRI experiment. Neuroimage. 2014;96:12–21.

40. Huang SC, Yu DC, Barrio JR, Grafton S, Melega WP, Hoffman JM, et al. Kinetics and modeling of L-6-[18F]fluoro-dopa in human positron emission tomographic studies. J Cereb Blood Flow Metab. 1991;11:898–913.

41. Luo WL, Nichols TE. Diagnosis and exploration of massively univariate neuroimaging models. Neuroimage. 2003;19:1014–32.

42. Becker JB, Chartoff E. Sex differences in neural mechanisms mediating reward and addiction. Neuropsychopharmacology. 2019;44:166–83.

43. Orsini CA, Willis ML, Gilbert RJ, Bizon JL, Setlow B. Sex differences in a rat model of risky decision making. Behav Neurosci. 2016;130:50–61.

44. Soutschek A, Burke CJ, Raja Beharelle A, Schreiber R, Weber SC, Karipidis, II, et al. The dopaminergic reward system underpins gender differences in social preferences. Nat Hum Behav. 2017;1:819–27.

45. Norris DG. Principles of magnetic resonance assessment of brain function. J Magn Reson Imaging. 2006;23:794–807.

46. Logothetis NK, Pauls J, Augath M, Trinath T, Oeltermann A. Neurophysiological investigation of the basis of the fMRI signal. Nature. 2001;412:150–7.

47. Krimer LS, Muly EC, 3rd, Williams GV, Goldman-Rakic PS. Dopaminergic regulation of cerebral cortical microcirculation. Nat Neurosci. 1998;1:286–9.

48. Knutson B, Gibbs SE. Linking nucleus accumbens dopamine and blood oxygenation. Psychopharmacology (Berl). 2007;191:813–22.

49. Sander CY, Hooker JM, Catana C, Normandin MD, Alpert NM, Knudsen GM, et al. Neurovascular coupling to D2/D3 dopamine receptor occupancy using simultaneous PET/functional MRI. Proc Natl Acad Sci U S A. 2013;110:11169–74.

50. Kehr W, Speckenbach W, Zimmermann R. Interaction of haloperidol and gamma-butyrolactone with (+)-amphetamine-induced changes in monoamine synthesis and metabolism in rat brain. J Neural Transm. 1977;40:129–47.

51. Hartvig P, Torstenson R, Tedroff J, Watanabe Y, Fasth KJ, Bjurling P, et al. Amphetamine effects on dopamine release and synthesis rate studied in the Rhesus monkey brain by positron emission tomography. J Neural Transm (Vienna). 1997;104:329–39.

52. Berry AS, Shah VD, Furman DJ, White RL, 3rd, Baker SL, O’Neil JP, et al. Dopamine Synthesis Capacity is Associated with D2/3 Receptor Binding but Not Dopamine Release. Neuropsychopharmacology. 2018;43:1201–11.

## SUPPLEMENTARY REFERENCES

1. Pessiglione M, Seymour B, Flandin G, Dolan RJ, Frith CD. Dopamine-dependent prediction errors underpin reward-seeking behaviour in humans. Nature. 2006;442:1042–5.

2. Rischka L, Gryglewski G, Pfaff S, Vanicek T, Hienert M, Klobl M, et al. Reduced task durations in functional PET imaging with [(18)F]FDG approaching that of functional MRI. Neuroimage. 2018;181:323–30.

3. Huang SC, Yu DC, Barrio JR, Grafton S, Melega WP, Hoffman JM, et al. Kinetics and modeling of L-6-[18F]fluoro-dopa in human positron emission tomographic studies. J Cereb Blood Flow Metab. 1991;11:898–913.

4. Doudet DJ, Chan GL, Jivan S, DeJesus OT, McGeer EG, English C, et al. Evaluation of dopaminergic presynaptic integrity: 6-[18F]fluoro-L-dopa versus 6-[18F]fluoro-L-m-tyrosine. J Cereb Blood Flow Metab. 1999;19:278–87.

